# Electron tomography analysis of the prolamellar body and its transformation into grana thylakoids in the cryofixed *Arabidopsis* cotyledon

**DOI:** 10.1101/2022.04.04.487035

**Authors:** Zizhen Liang, Wai-Tsun Yeung, Juncai Ma, Keith Ka Ki Mai, Zhongyuan Liu, Yau-Lun Felix Chong, Xiaohao Cai, Byung-Ho Kang

**Author notes:** Address correspondence to or. The author responsible for distribution of materials integral to the findings presented in this article in accordance with the policy described in the Instructions for Authors (www.plantcell.org) is: Byung-Ho Kang.

## Abstract

The para-crystalline structure of prolamellar bodies (PLBs) and light-induced etioplasts-to-chloroplasts transformation have been investigated with electron microscopy methods. However, these studies suffer from chemical fixation artifacts and limited volumes of three-dimensional reconstruction. We have examined Arabidopsis thaliana cotyledon cells with electron tomography (ET) to visualize etioplasts and their conversion into chloroplasts. We employed the scanning mode of ET for imaging large volumes and high-pressure freezing to improve sample preservation. PLB tubules were arranged in a zinc blende-type lattice like carbon atoms in diamonds. Within 2 hours after illumination, the lattice collapsed from the PLB exterior and the disorganized tubules merged to form thylakoid sheets, a.k.a. pre-granal thylakoids. These pre-granal thylakoids in PLB’s vicinity folded and overlapped with each other to create grana stacks. Since the nascent pre-granal thylakoids had curved membranes in their tips, we examined the expression and localization of CURT1 proteins. *CURT1A* transcript was most abundant in de-etiolating cotyledon samples, and CURT1A concentrated to the PLB periphery. In *curt1a* etioplasts, PLB-associated thylakoids were swollen and failed to form grana stacks. By contrast, PLBs had cracks in their lattices in *curt1c* etioplasts. Our data provide evidence that CURT1A is required for pre-granal thylakoid assembly from PLB tubules during de-etiolation, while CURT1C contributes to the cubic crystal growth under darkness.

## Introduction

Plastids exist in different forms depending on the cell type and environmental conditions (Jarvis and López-Juez, 2013). In germinating seedlings, proplastids in the cotyledon develop into chloroplasts. When chlorophyll biosynthesis is inhibited in the absence of light, the photosynthetic protein complexes of the thylakoid membrane are not assembled, and chloroplast biogenesis is inhibited (Leivar et al., 2008). Instead, developmentally arrested plastids, known as etioplasts, form (Solymosi and Schoefs, 2010). Etioplasts transform into chloroplasts once light becomes available and chlorophyll accumulates (Hernandez-Verdeja et al., 2020).

Thylakoids in etioplasts consist of semi-crystalline tubular membrane networks of prolamellar bodies (PLBs) connected by planar prothylakoids (Ryberg and Sundqvist, 1982; Rascio et al., 1984). During the light-induced etioplast-chloroplast transition, lipids and cofactors stored in PLBs provide building blocks for the chloroplast thylakoids (Ploscher et al., 2011; Armarego-Marriott et al., 2019; Fujii et al., 2019). The most abundant protein constituent of PLBs is light-dependent protochlorophyllide oxidoreductase (LPOR) (Blomqvist et al., 2008), which forms a helical array surrounding PLB tubules (Floris and Kuhlbrandt, 2021). LPOR is a photocatalytic enzyme that mediates the reduction of protochlorophyllide (Pchlide) into chlorophyllide to produce chlorophyll (Zhang et al., 2019). LPOR oligomerizes on liposomes mimicking the PLB membrane to tubulate them *in vitro* as shown by cryo-electron microscopy (Nguyen et al., 2021). It is thought that LPOR undergoes conformational changes after the photoreduction and dissociates from the PLB membrane, resulting in the breakdown of the PLB lattice. Inactivation of *PORA*, an *Arabidopsis* gene encoding an LPOR protein, led to structural defects in PLBs and abnormal photomorphogenesis (Paddock et al., 2012).

When examined under electron microscopy, PLBs are made of hexagonal lattices in which tetrahedral units repeat (Murakami et al., 1985). Small angle X-ray studies of isolated PLBs revealed that branched tubules in PLBs are packed primarily in the cubic diamond (*i.e*., zinc blende) symmetry (Williams et al., 1998; Selstam et al., 2007). Recent electron tomography (ET) imaging of runner bean (*Phaseolus coccineus*) indicated that the PLB lattice matched the wurtzite-type crystal symmetry (Kowalewska et al. 2016). PLBs in which tubules deviate from the tetrahedral pattern have been reported, and the arrangement is called the “open” type (Gunning, 2001). Moreover, etiolation conditions affect the sizes and density of PLBs (Bykowski et al., 2020).

The light-triggered transformation of PLBs into grana and stroma thylakoids was first investigated with electron microscopy in the 1960s, although those early studies were based on two-dimensional electron micrographs of PLBs and thylakoids (Gunning, 1965; Henningsen and Boynton, 1974; Rascio et al., 1984; Grzyb et al., 2013). An ET analysis of the PLB and thylakoids in de-etiolating runner bean cotyledons showed that PLB tubules directly change into planar thylakoid elements without the involvement of vesicular intermediates and that the helical arrangement of the inter-disc connections within a grana stack appears early in the granum development (Kowalewska et al., 2016). In the ET study, the etioplast volumes in the 3D reconstruction were limited in the z-direction coverage, visualizing two hexagonal layers with a 120 kV electron microscope. Using serial block-face scanning electron microscopy, it was demonstrated that the conversion of PLBs into photosynthetic thylakoids in *Arabidopsis* cotyledons occurs within 24 hour after illumination in concurrence with correlative proteomic and lipidomic results (Pipitone et al., 2021). The serial block-face scanning electron microscopy approach can visualize larger volumes encompassing the entire etioplasts or chloroplasts, but the resolution is poorer than ET, especially along the z-axis. In both 3D EM studies, cotyledon samples were prepared with chemical fixation, which fails to preserve intricate or short-lived structures in cells (McIntosh et al., 2005; Staehelin and Kang, 2008).

In this study, we examined etioplasts in *Arabidopsis* cotyledons grown in the dark with serial section ET. Cotyledon samples were prepared by high-pressure freezing to avoid fixation artifacts. As the stroma is heavily stained in high-pressure frozen etioplasts, we employed scanning transmission ET (STET), which enhances image contrast in tomograms from such specimens (Aoyama et al., 2008; Hohmann-Marriott et al., 2009; Murata et al., 2014; Kang, 2016). CURVATURE THYLAKOID1 (CURT1) family proteins are thylakoid membrane proteins that stabilize the sharply curved membrane at the grana margin (Armbruster et al., 2013; Pribil et al., 2014). Our time-resolved ET study of wild-type and *curt1* family mutant cotyledons indicate that grana stacks arise directly from PLBs and that CURT1A is required for the stack assembly. By contrast, CURT1C plays a role in the cubic crystal close packing for PLB biogenesis in developing etioplasts.

## Results

### Crystalline structure of the *Arabidopsis* PLBs and their light-induced degradation

To estimate the timeline of the etioplast-to-chloroplast transformation, we examined etiolated *Arabidopsis Col-0* cotyledons at 0, 1, 2, 4, 8, and 12 hours after illumination (HAL). Cotyledon greening was clearly noticed at 12 HAL (Fig. S1 A-C), and chlorophyll autofluorescence increased during this period (Fig. S1 D-H). PLBs in 0 HAL cotyledons were small spots of 1.5-2.0 μm in diameter that emitted autofluorescence (Fig. S1 D and D’). The fluorescent spots enlarged and spread at 2 HAL, as chlorophyll molecules were produced from Pchlide in PLBs and mobilized (Fig. S1 E and E’). We monitored the degradation of PLBs in high-pressure frozen cotyledon samples with transmission electron microscopy (TEM) and STET at the six time points. Each etioplast had PLBs and prothylakoids radiating from PLBs at 0 HAL (Fig. 1A). These prothylakoids were planar and had ribosomes on their stroma surface (Fig. S1 J-K), resembling the pre-granal thylakoids of proplastids in germinating cotyledon cells at 36 and 64 hours after imbibition (Liang et al. 2018).

**Figure 1.**
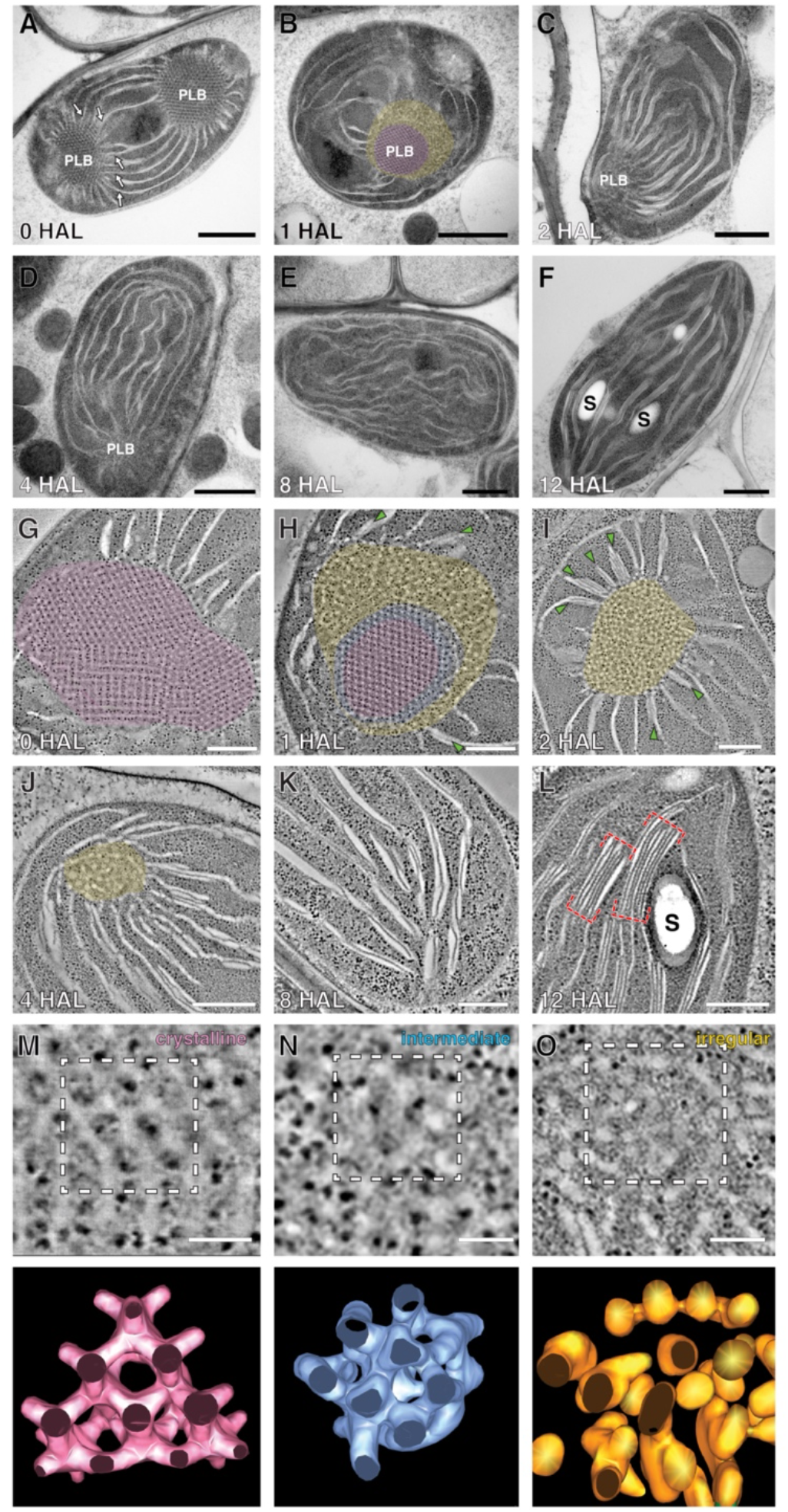
PLB degeneration and thylakoid assembly in de-etiolating Arabidopsis cotyledons. (A-F) TEM micrographs of etioplasts/chloroplasts at A) 0 HAL, B) 1 HAL, C) 2 HAL, D) 4 HAL, E) 8 HAL, and F) 12 HAL. Arrows in (A) indicate prothylakoids. S: starch particle. Scale bars = 1 μm. (G-L) STET slice images of plastids at G) 0 HAL, H) 1 HAL, I) 2 HAL, J) 4 HAL, K) 8 HAL, and L) 12 HAL. The crystalline, irregular, and intermediate zones in PLBs are highlighted in magenta, yellow, and blue, respectively, in (B) and (G-J). The intermediate zone was distinguished by STET (H) but not by TEM (B). PLB-associated grana stacks are marked with green arrows in (H) and (I). Grana stacks are denoted with red brackets in (L). S: starch particle. Scale bars = 300 nm. (M-O) High magnification STET slice images of the PLB lattice (crystalline) at 0 HAL (M), PLB tubules of the intermediate zone at 1 HAL (N), and PLB tubules of the irregular zone at 1 HAL (O). Scale bars = 150 nm. Lower panels show 3D surface models of the PLB membranes demarcated with dashed squares in upper images.

PLBs shrank quickly and lost their crystalline regularity by 2 HAL (Fig. 1 A-C, G-I). Double layered thylakoids appeared in PLBs’ vicinity as early as 1 HAL (Fig. 1H), and the number of disks increased in the PLB-associated grana stacks at 2 HAL (Fig. 1I). PLBs almost degraded in 4 HAL samples and disappeared completely by 8 HAL (Fig. 1D-E and J-K). Chloroplasts at 12 HAL had typical thylakoid networks where grana stacks consisted of as many as 6-8 disks and they were interconnected by stroma thylakoids (Fig. 1 F and L). They had starch particles and were approximately 25% larger than etioplasts at 0 HAL (Fig. 1L and S1I).

Loss of the crystalline architecture began from the PLB surface (Fig. 1B). The inner core retained the lattice structure in 1 HAL PLBs, while tubules at the periphery became disorganized (Fig. 1 B and H). Between the crystalline core and the irregular periphery lied a narrow band in which the lattice arrangement was compromised when examined with STET (Fig. 1H). No crystalline symmetry was discerned in PLBs at 2 HAL. Many grana stacks arose in association with PLBs (Fig. 1I), suggesting that chlorophyll molecules produced from protochlorophyllides in the PLB are directly incorporated into photosystem II (PSII) – light harvesting complex II (LHCII) supercomplexes that concentrate to the appressed thylakoid regions in the grana stack (Daum et al., 2010; Wietrzynski et al., 2020)

We generated 3D surface models of PLB tubules in the crystalline core, disorganized periphery, and the intermediate zones at 0 and 1 HAL. PLBs before illumination consisted of tetravalent nodes (Fig. 1M), and the average width of the tubules was calculated to be 24.0 nm (Fig. 2K). In the intermediate zone at 1 HAL, tubular nodes were displaced, obscuring the hexagonal pattern (Fig. 1N). The peripheral tubules were highly convoluted with varying thicknesses (Fig. 1O).

**Figure 2.**
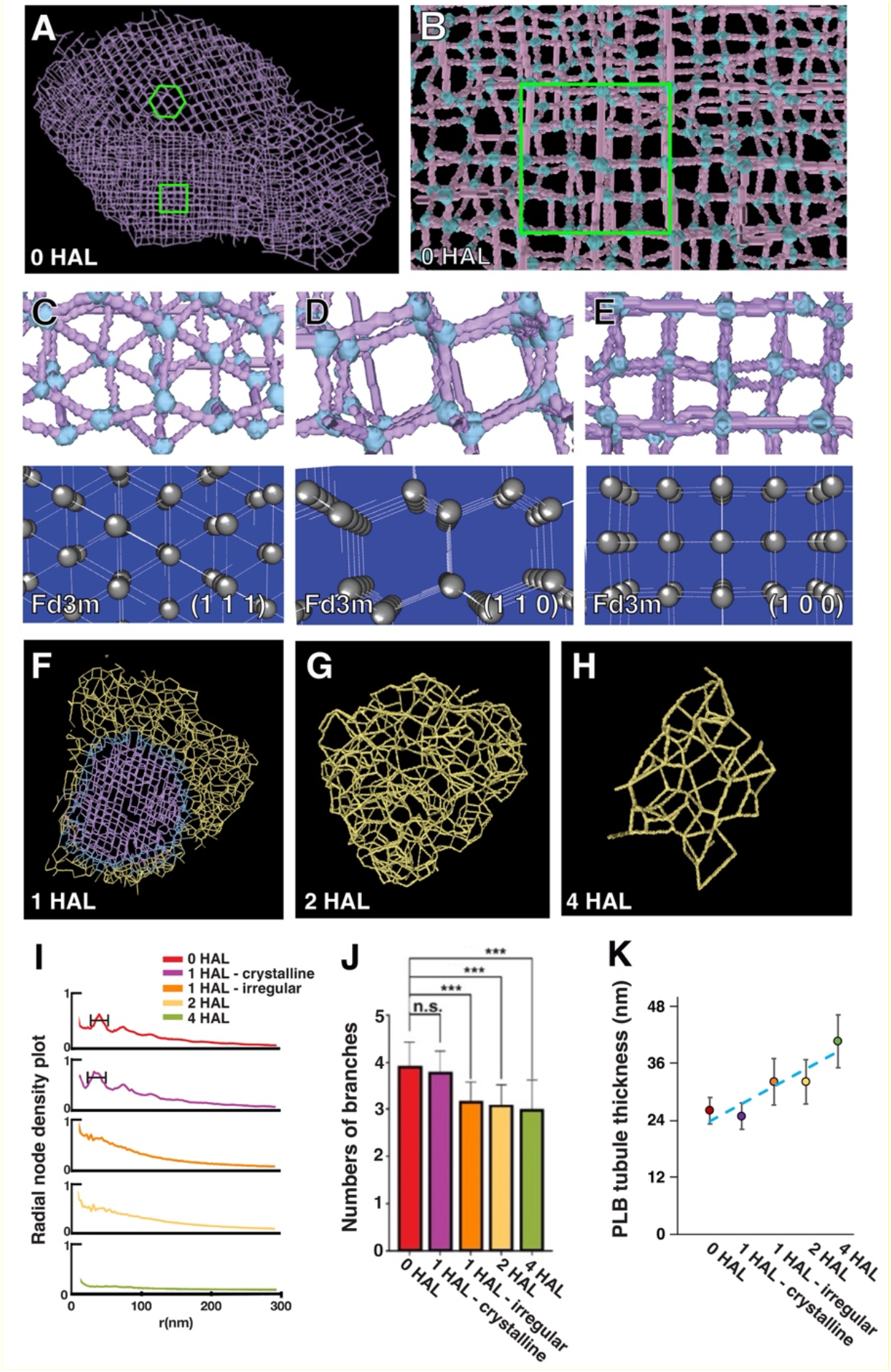
The crystalline structure of Arabidopsis PLB and its decay during de-etiolation. (A) A skeleton model of the PLB in Fig. 1G. Regions exhibiting hexagonal or square lattice patterns are marked in green. (B) A higher magnification view of the skeleton model shown in panel A. Nodes are highlighted in light blue. The region exhibiting a square lattice pattern is marked with a green square. (C-E) Projection views of select regions (upper panels), and lattice planes of the space group Fd3m (cubic diamond crystal structure) and their Miller indices, (1,1,1), (1,1,0), and (1,0,0) of the PLB skeleton model (bottom panels). Note that arrangements of PLB nodes and tubules match those of the cubic diamond lattices in all three planes. (F-H) Skeleton models of decaying PLBs at F) 1 HAL, G) 2 HAL, and H) 4 HAL. The models were generated from the tomograms in Fig. 1H, I, and J, respectively. Lines are color-coded to denote the crystalline, irregular, and intermediate zones in PLBs. (I) Radial density plots of branching nodes at four timepoints of de-etiolation. (J) The average numbers of branches at each node in 0 HAL, 1 HAL crystalline, 1 HAL irregular, 2 HAL, and 4 HAL PLBs. Branches were counted from 24 nodes at each stage. (***: p-value<0.0005 by Welch’s t-test, n.s., no significant difference) (K) The average thicknesses of tubules in 0 HAL, 1 HAL crystalline, 1 HAL irregular, 2 HAL, and 4 HAL PLBs. The thicknesses were calculated from 81225 (0 HAL), 17343 (1 HAL crystalline), 20864 (1 HAL irregular), 13391 (2 HAL), and 2333 (4 HAL) tubular segments in PLB surface models.

### 3D skeleton models of PLBs

To determine the crystalline symmetry of PLBs and analyze their collapse quantitatively, we created 3D skeleton models of PLBs in our STET tomograms (Fig. S2). In the models consisting of lines and nodes, hexagonal and square lattices were readily discerned in PLBs at 0 HAL (Fig. 2A-B). We were able to capture projection views from the models matching the Miller indices of the diamond cubic symmetry (Fig. 2C-E). The periodic hexagonal patterns conformed to the (1,1,1) or (1,1,0) planes, whereas the square lattice matched the (1,0,0) plane. The diamond cubic unit cell size averaged to 65.5 nm (n = 61, SD = 3.71 nm) when measured from nodes in the (1,1,0) or (1,0,0) planes.

From the skeleton models from 1, 2, and 4 HAL PLBs (Fig. 2F-H), we calculated radial densities of nodes and branching numbers per node at each time point. The node density plot had a peak from 30 nm to 70 nm in 0 HAL and 1 HAL crystalline PLBs, indicating a regular spacing between nodes (Fig. 2I). The peak was not present in the skeleton model from the irregular region of PLBs at 1 HAL. In agreement with the tetravalent units seen in 3D models (Fig. 1M), each node had four branches in 0 HAL and 1 HAL crystalline PLBs (Fig. 2J). The numbers of branches decreased as PLBs were degraded in later time points (Fig. 2J) and the reduction was accompanied by an increase in tubule thickness (Fig. 2K).

### Assembly of pre-granal thylakoids and grana stacks on the PLB surface

3D tomographic models of the PLB-thylakoid interface at 1, 2, and 4 HAL were generated to examine how PLB tubules give rise to pre-granal thylakoids and how they turn into grana stacks. The irregular tubules observed in the PLB periphery became interwoven and smoothened to constitute fenestrated membrane sheets at 1 and 2 HAL (Fig. 3A-E). Thylakoids in the immediate vicinity of PLBs were fenestrated but they consolidated into pre-granal thylakoids. Fenestrae shrank and disappeared by 400 nm away from PLBs (Fig. 3L).

**Figure 3.**
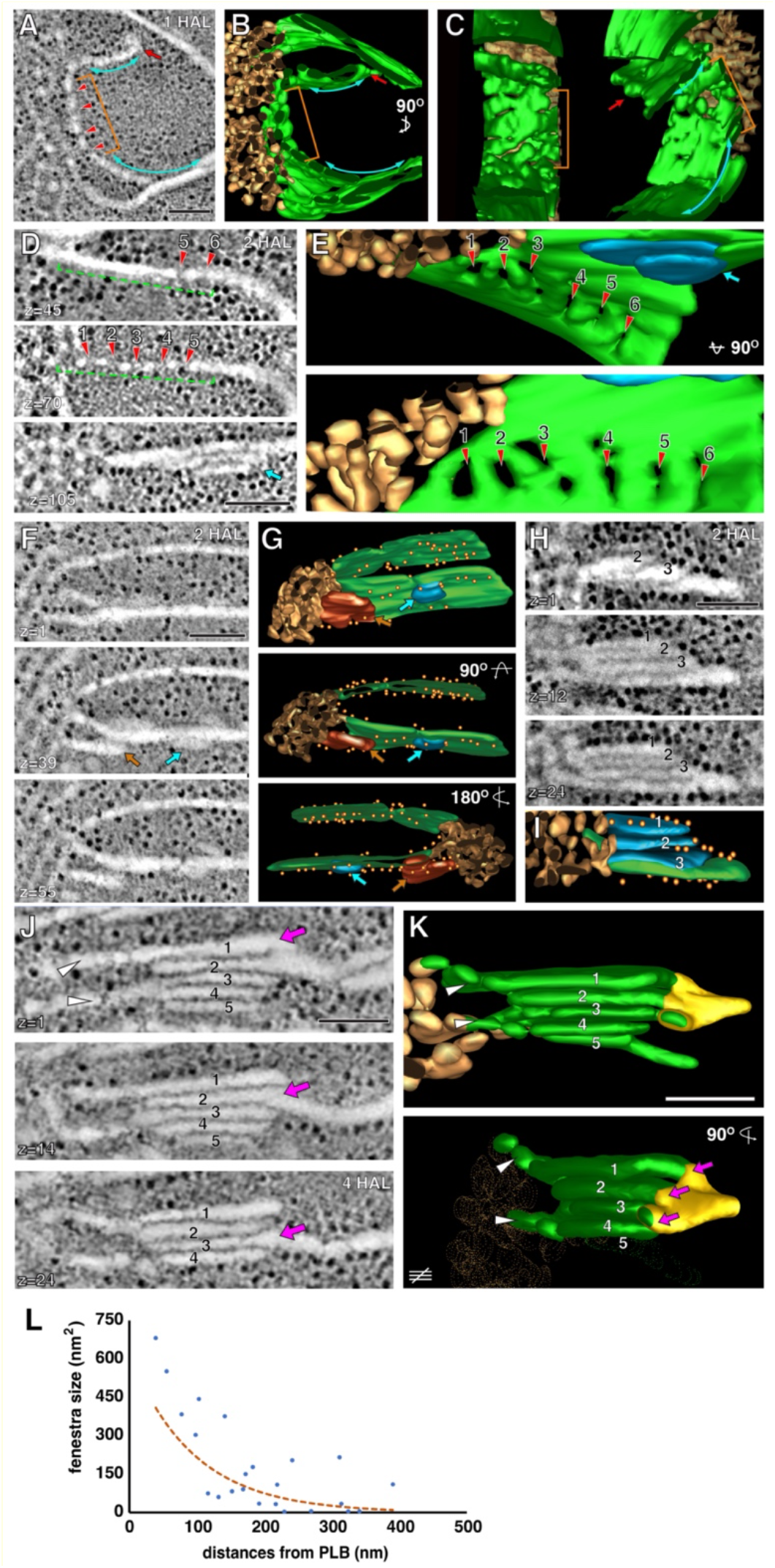
PLB to pre-granal thylakoid transition and grana formation from pre-granal thylakoids. (A-C) STET slice image (A) and 3D models (B-C) of a PLB (gold) and prothylakoids (blue double-sided arrows) and a fenestrated sheet between them (brown bracket) at 1 HAL (red arrows in A-C) (D-E) Fenestrate sheet connected to a PLB at 2 HAL. Fenestrae are indicated with red arrowheads. (F-G) High-magnification images of two pre-granal thylakoids connected to a PLB at 2 HAL. A bud emerging from the pre-granal thylakoid is marked with a blue arrow. (H-I) Image and 3D model of a nascent granum consisting of four layers at the margin of a PLB at 2 HAL. Three disks (blue) derived from the irregular tubules pile up on a grana-forming thylakoid (green). They are interconnected via their margin. (J-K) STET slice image (J) and 3D models (K) of a granum and stroma thylakoids associated with a PLB at 4 HAL. The granum consists of five disks that are linked via a helical thylakoid arrangement (yellow membrane in K). As the slice number increases from 1 to 21, the disks 1, 2, and 3 make connections sequentially to the stroma thylakoid (magenta arrows in J and K). Scale bars = 100 nm. (L) Correlation plot illustrating the relation between fenestrae sizes and their distances from PLBs at 2 HAL.

Stacked thylakoids developed from pre-granal thylakoids in the immediate vicinity of PLBs at 2 HAL (Fig. 3F-G). Pre-granal thylakoids laterally overlapped (brown arrow in Fig. 3G) or tongue-like outgrowths emerged from and lay down over pre-granal thylakoids (blue arrow in Fig. 3G). Three or four-layered grana appeared where thylakoids repeatedly folded (Fig. 3H-I). The acquisition of new layers did not seem to occur in an orderly fashion. Their diverse membrane configurations were similar to those of pro-granal stacks in young chloroplasts of germinating cotyledon cells (Liang et al. 2018). Grana stacks displaced from PLBs were frequently observed at 4 HAL (Fig. 3J-K). They had five or six disks linked by helical stroma thylakoids as a typical granum does.

We performed RNA-seq and immunoblot analysis of de-etiolating cotyledon samples isolated at 0, 1, 2, 4, 8, and 12 HAL (Fig. S3). Most components of the photosystems and light-harvesting complexes were upregulated at 2 HAL when grana stacks appeared on the PLB surface (Fig. S3A). However, polypeptide levels of PSII, LHCII, PSI, and LHCI did not exhibit a significant increase until 8 HAL, when PLB disappeared except for PsaD2 (Fig. S3 B-C). The PORA polypeptide levels gradually decreased soon after light exposure and its transcript amounts were low except for 0 HAL samples (Fig. S3 A-C).

### CURT1A localized to the nascent pre-granal thylakoids emerging from PLBs and grana stacks

Pre-granal thylakoids and grana stacks emerging from the PLB surface had highly bent membranes exposed to the stroma (Fig. 3A-C, red arrows). This led us to hypothesize that CURT1 family proteins, which stabilize the curved membrane of each disk in the grana stack, are involved in the grana assembly from PLB tubules (Armbruster et al., 2013; Pribil et al., 2014). In the RNA-seq dataset from de-etiolating Arabidopsis cotyledon samples, mRNA levels of *CURT1A* (AT4G01150), *CURT1B* (AT2G46820), and *CURT1C* (AT1G52220) were low at 0 HAL. CURT1A transcription was more active among the three members, with its transcript amounts increased by about six-fold (Fig. S3 A and B). In immunoblot analysis, CURT1A, 1B, and 1C were detected in 0 HAL samples, indicating that they accumulate in PLBs during skotomorphogenesis (Fig. S3 C and D). Amounts of the CURT1A polypeptide approximately doubled, while the CURT1B and CURT1C polypeptide levels did not change. The smaller changes in the CURT1 polypeptide amounts than those of *CURT1* transcripts suggest that turnover of CURT1 proteins occurs in de-etiolating cotyledon cells.

We generated transgenic *Arabidopsis* lines expressing a CURT1A-GFP fusion protein under control of its native promoter to monitor its localization. The fusion protein rescued the granum assembly defects of *curt1a-1* mutant cotyledons, indicating that the fusion protein is functional (Fig. S4 and S5). At 0 HAL, GFP fluorescence partially overlapped with PLB autofluorescence; some PLBs had a GFP halo or GFP-positive puncta around them (Fig. 4A). CURT1A-GFP formed foci on PLBs in 2 HAL chloroplasts over which chlorophyll autofluorescence spread (Fig. 4B). Small GFP spots scattered to multiple locations that could correspond to where grana stacks develop in 4 HAL and 8 HAL chloroplasts (Fig. 4 C and D). We verified the localization of CURT1A with PLBs with immunogold labeling at the four time points. CURT1A-specific gold particles were associated most frequently with the PLB cortices where new pre-granal thylakoids assembled at 0 to 2 HAL (Fig. 4 E-I, brown arrowheads, and M). As PLBs shrank at 4 and 8 HAL, the majority of CURT1A relocated to thylakoids, binding to grana margins (Fig. 4J-L, green arrowhead, and M). PLBs had CURT1A-GFP and CURT1A immunogold particles at 0 HAL, validating that CURT1A is deposited in PLBs under darkness. This observation agrees with the discrepancy between mRNA and polypeptide levels of CURT1A at 0 HAL (Fig. S3)

**Figure 4.**
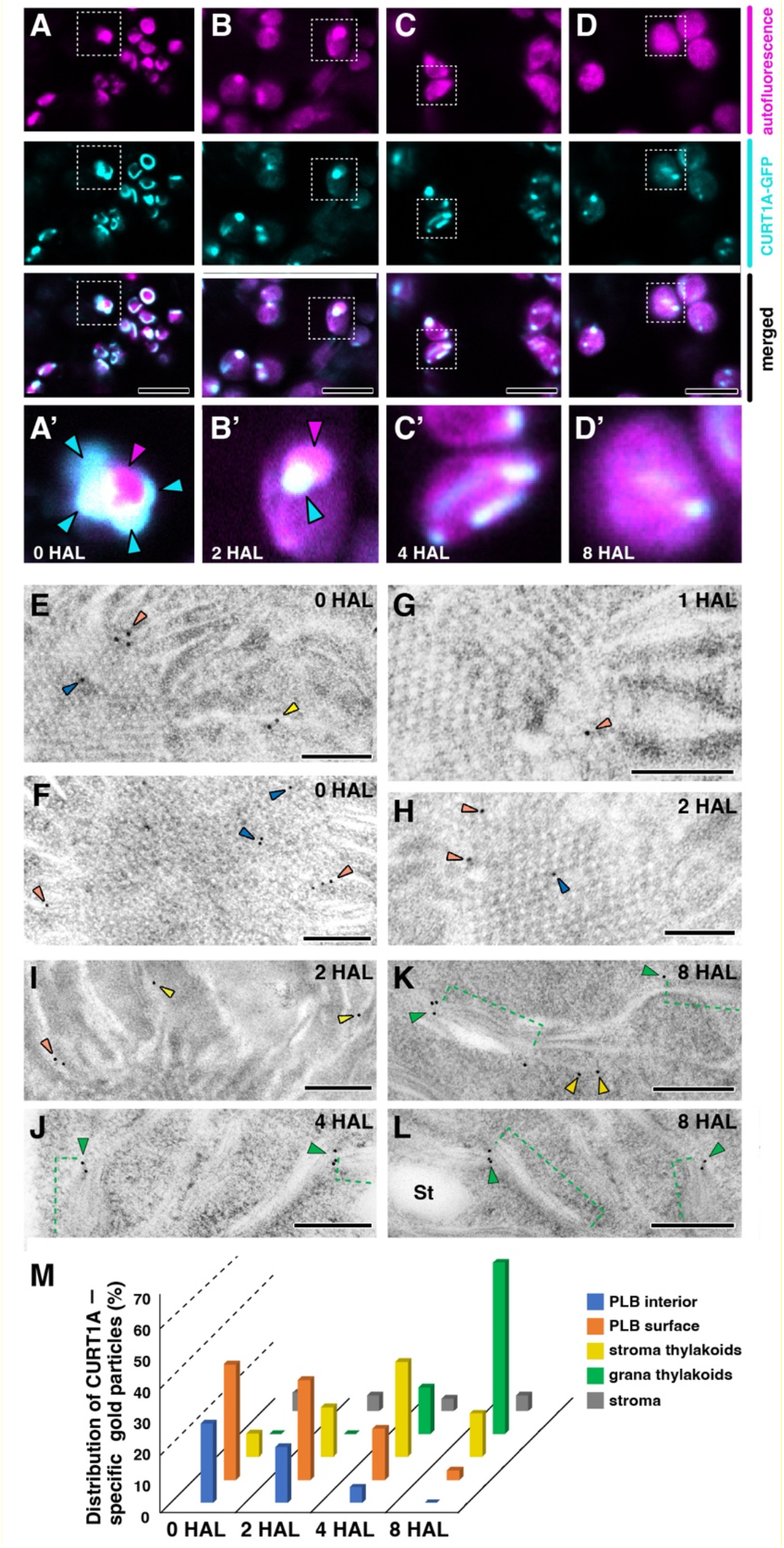
Localization of CURT1A in PLBs and thylakoids. (A-D) Confocal laser scanning micrographs showing CURT1A-GFP distribution at A) 0 HAL, B) 2 HAL, C) 4 HAL, and D) 8 HAL. Autofluorescence from Pchlide/chlorophyll, fluorescence from CURT1A-GFP, and merged panels are shown in each column. Panels A’-D’ are high-magnification micrographs of regions indicated with squares in panels A-D. In A’ and B’, PLBs and CURT1A-GFP puncta are indicated with magenta and blue arrowheads. Scale bars = 8 μm. (E-L) Immunogold labeling localization of CURT1A in *Arabidopsis* plastids at E-F) 0 HAL, G-H) 1 HAL, I) 2 HAL, J) 4 HAL, and K-L) 8 HAL. Gold particles located in PLBs, periphery of PLBs, stroma thylakoids, and grana stacks (green brackets in J-L) are marked with blue, orange, yellow, and green arrowheads, respectively. Scale bars = 200 nm. (M) Histogram showing CURT1A-specific gold particle distribution in *Arabidopsis* plastids at 0 HAL, 2 HAL, 4 HAL, and 8 HAL. 200-300 gold particles were counted in 30-40 TEM sections from three different cotyledon samples at each time points.

### Aberrant assembly of pre-granal thylakoids and grana stacks in *curt1a* etioplasts

To test whether CURT1A is required for the pre-granal thylakoid assembly and grana formation, we isolated *curt1a* T-DNA inserted mutant lines (Fig. S4 and S5). Etioplasts in 0 HAL *curt1a-1* (SALK_030000) cotyledon cells appeared normal, and they had crystalline PLBs (Fig. 5A). We were able to capture projection views matching the cubic diamond crystal system (Fig. 5 B-C). However, at 0 and 1 HAL, thylakoids on the PLB surface were swollen in mutant cotyledon cells in contrast to the flat pre-granal thylakoids in wild-type cotyledon cells (Fig. 5 A-D). The swollen thylakoids between a PLB and prothylakoids had fenestrae but they failed to form grana stacks. (Fig. 5 O-P).

**Figure 5.**
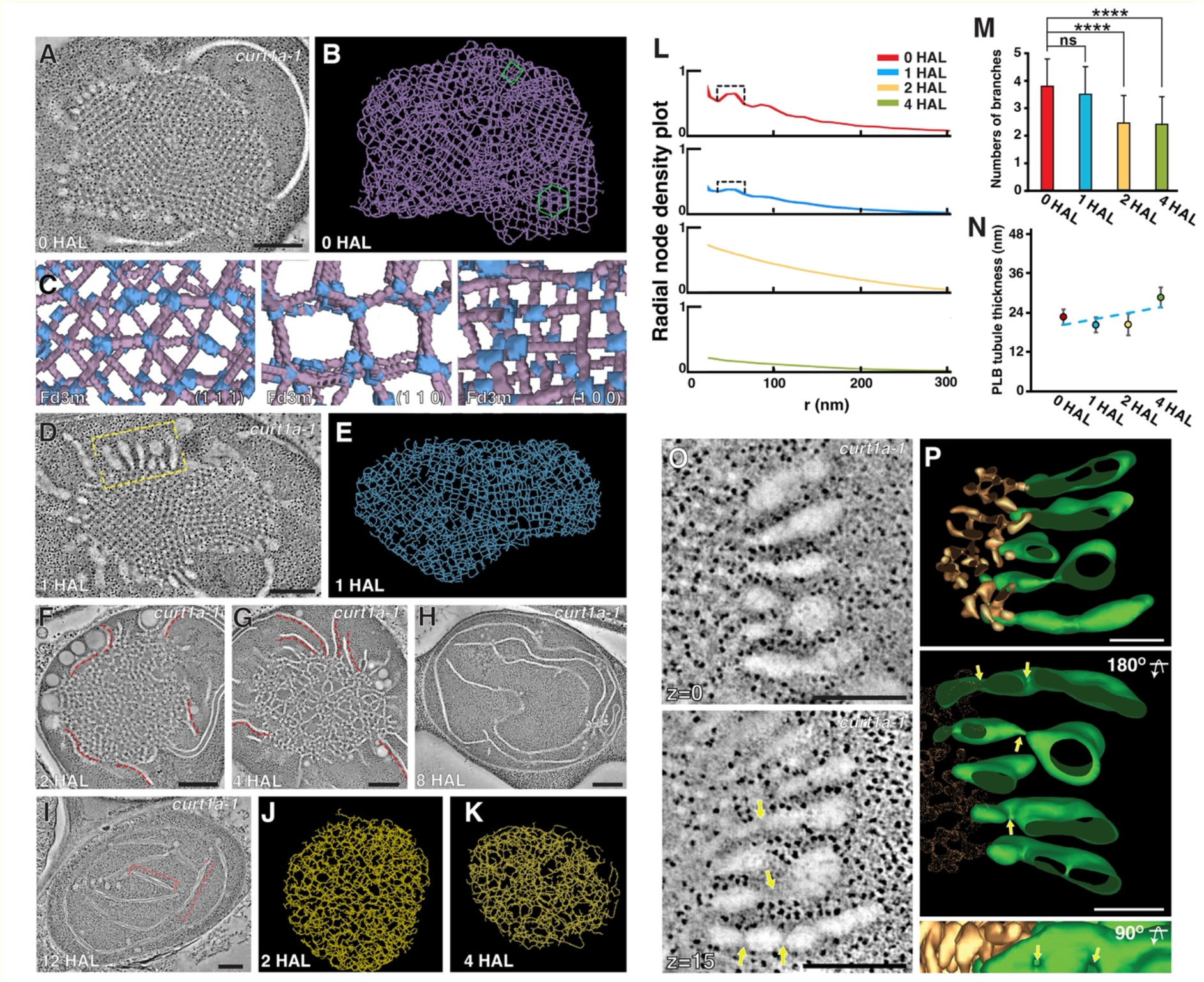
Altered PLB-to-thylakoid conversion in the *curt1a* etioplast. (A-B) STET slice image (A) of a *curt1a-1* plastid at 0 HAL and a skeleton model of its PLB (B). Regions with hexagonal or square lattice patterns are marked with a green hexagon or square, respectively, in (B). (C) Projection views of select regions in the skeleton model matching the space group Fd3m. Miller indices, (1,1,1), (1,1,0), and (1,0,0) of the lattice planes are indicated. (D-E) STEM tomography slice image (D) of a *curt1a-1* plastid at 1 HAL and a skeleton model of its PLB (E). (F-I) STET slice image of *curt1a-1* plastids at F) 2 HAL, G) 4 HAL, H) 8 HAL, and I) 12 HAL. Red dots in (F) and (G) label pre-granal thylakoids associated with PLBs. Note that grana stacks failed to form around PLBs. Scale bars in (A, D, F-I) = 300 (J-K) Skeleton models of decaying PLBs at J) 2 HAL and K) 4 HAL. (L) Radial density plots of branching nodes at the four timepoints of de-etiolation. Radial distances with higher node densities in 0 and 1 HAL plots are indicated with brackets. (M) The average numbers of branches at each node in 0 HAL, 1 HAL, 2 HAL, and 4 HAL PLBs. Branches were counted from 21 nodes at each stage. (****: p-value<0.0001 by Welch’s t-test, n.s., no significant difference) (N) The average thicknesses of tubules in 0 HAL, 1 HAL, 2 HAL, and 4 HAL PLBs. The thicknesses were calculated from 52306 (0 HAL), 10106 (1 HAL), 17058 (2 HAL), and 42453 (4 HAL) tubular segments in PLB surface models. (O) A tomographic slice image of the thylakoids connected to PLBs in the yellow bracket in (D). Fenestrations in pre-granal thylakoids are indicated with arrows. Scale bars = 200 nm. (P) 3D model of the swollen thylakoids (green) and PLB (gold) in (O). Four fenestrae marked with arrows in (P) correspond to the four in the lower panel on (O). Scale bars = 100 nm.

Unlike the 1 HAL PLB in wild-type that consisted of of the crystalline core and irregular periphery (Fig. 1H and 2F), PLBs in *curt1a-1* at 1 HAL retained the lattice architecture throughout its volume (Fig. 5D). In the skeleton model (Fig. 5E), nodes were separated by regular intervals (Fig. 5L, brackets) and their degree of branching did not change (Fig. 5M) in 1 HAL PLBs. *curt1a* PLBs did not exhibit any crystalline structure at 2 HAL as they shrank, and stroma thylakoids proliferated (Figs. 5 F, J, L-M). PLBs disappeared by 8 HAL but no grana stacks were found in *curt1a* chloroplasts (Figs. 5 G-H). Chloroplasts in *curt1a-1* mutant cotyledon cells at 12 HAL had extremely wide grana stacks made of two or three disks (Fig. 5I). Another T-DNA mutant allele of curt1a (*curt1a-2*, GK-805B04) also had crystalline PLBs, swollen thylakoids at 1 HAL, and lacked grana stacks (Fig. S5 A-E). The *curt1a-1* phenotypes were rescued by transformation with the CURT1A-GFP construct (Fig. S5 G-J).

### The cubic crystalline lattice was disrupted in *curt1c* PLBs

As transcripts from *CURT1B* (AT2G46820) and *CURT1C* (AT1G52220) accumulated in de-etiolating cotyledon cells, we examined T-DNA mutant lines in which *CURT1B* or *CURT1C* was inactivated (Figs. 6 and S6). We noticed that PLBs of *curt1c-1* (SALK_023574) cotyledons often had irregularities in their crystalline structure (Fig. 6 A-C, and I). Pores as large as 400 nm (Fig. 6A) or areas where PLB tubules were disorganized (Fig. 6B) were also seen in *curt1c-1* PLBs. However, PLBs in 1 HAL *curt1c-1* etio-chloroplasts had irregular tubules in the PLB periphery and a crystalline center (Fig. 6D). These observations contrasted with the PLB phenotypes of *curt1a-1*. (Fig. 5 D-E)._In 2 and 4 HAL mutants, grana stacks developed in association with degrading PLBs (Fig. 6E-F). When we expressed CURT1C-GFP from its native promoter in the *curt1c-1* background, the defects in PLBs disappeared (Fig. 6 H and I).

**Figure 6.**
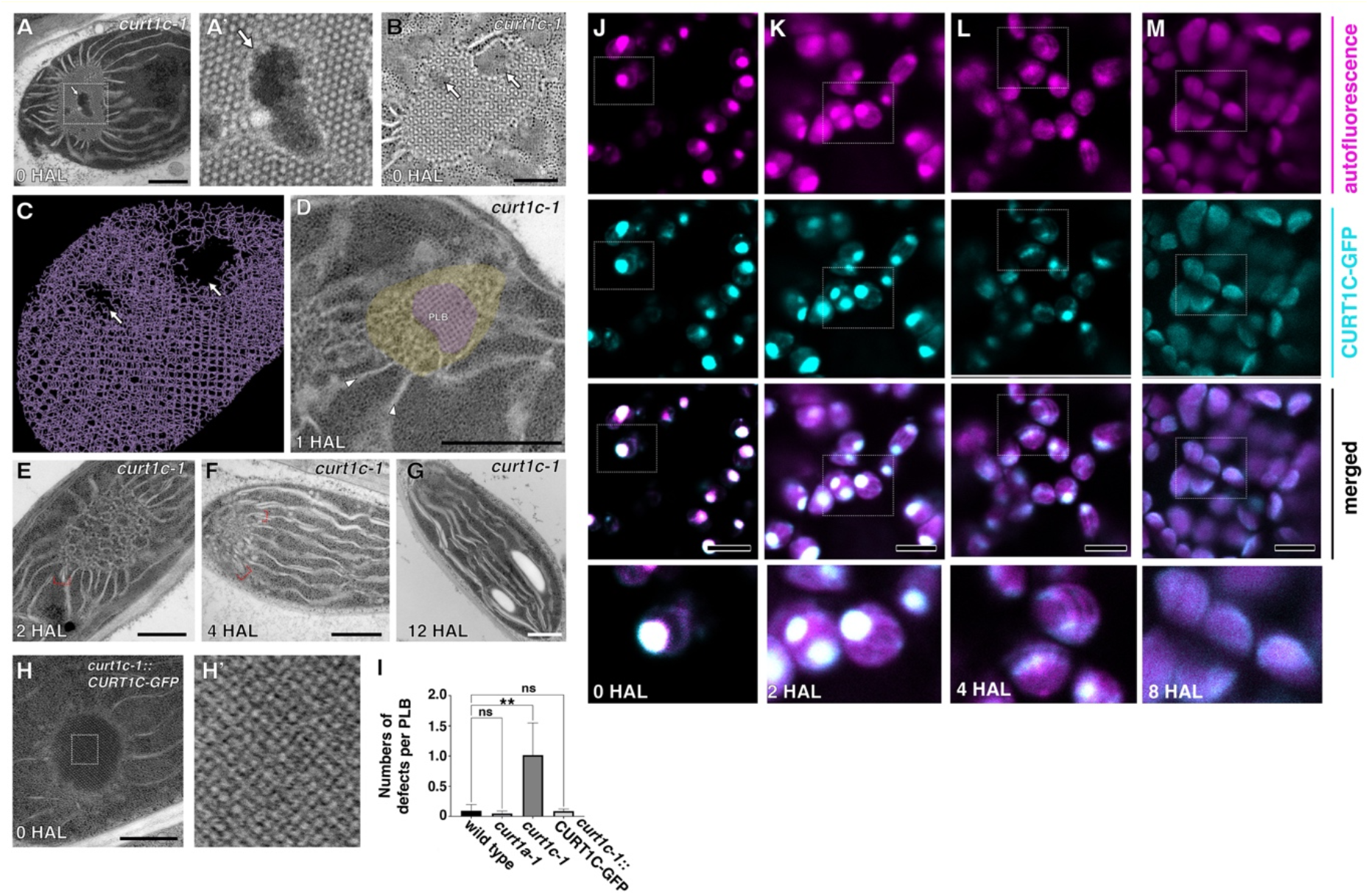
PLBs in *curt1c* mutant etioplasts are abnormal. (A-B) TEM micrographs (A) and STET slice image (B) of PLBs in *curt1c-1* plastids at 0 HAL. Arrows mark defects in the PLB lattice. (C) A skeleton model of *curt1c-1* PLB in (B). Arrows point to the pores in the PLB. (D) TEM micrograph of a *curt1c-1* plastid at 1 HAL. Zones in the PLB with irregular and crystalline tubules are highlighted in yellow and magenta, respectively. (E-G) TEM micrographs of *curt1c-1* plastids at 2 HAL (E), 4 HAL (F), and 12 HAL (G). Grana stacks associated with PLBs are denoted with red brackets in (E) and (F). Scale bars = 500 nm. (I) Histogram showing pore numbers per etioplast. ~30 etioplasts in TEM sections from at least three samples for each genotype were examined. (±SD; one-way ANOVA; **p < 0.01, n.s., no significant difference). (H) TEM micrograph of an etioplast in *curt1c-1* expressing CURT1C-GFP at 0 HAL. (A’) and (H’) are magnified views of PLBs inside the rectangles in (A) and (H), respectively. (J-M) CURT1C-GFP localization at 0 HAL (J), 2 HAL (K), 4 HAL (L), and 8 HAL (M) visualized by confocal laser scanning microscopy. Autofluorescence from Pchlide/chlorophyll, fluorescence from CURT1C-GFP, merged panels, and higher-magnification micrographs of regions denoted with squares are provided in each column. Scale bars in (J-M): 8 μm.

CURT1C-GFP expressed by the *CURT1C* promoter overlapped almost entirely with PLB autofluorescence at 0 HAL and shrank together with PLBs (Fig. 6 J-M). In 4 HAL etio-chloroplasts where PLBs have been mostly depleted, GFP-positive spots were scattered over thylakoids (Fig. 6L). At 8 HAL when PLBs were completely disappeared, small GFP spots were no longer distinguished. Instead, CURT1C-GFP seemed to spread over the entire chloroplast (Fig. 6M). PLBs, their degradation, and thylakoid development around PLBs appeared normal in *curt1b-1* (WiscDsLoxHs047_09D) cotyledon cells (Figs. S6).

## Discussion

We have determined the crystalline structure of *Arabidopsis* PLBs in high-pressure frozen etioplast samples to be zinc blende type. This result agrees with a small angle X-ray diffraction study of isolated maize PLBs (Selstam et al., 2007). Floris and Kühlbrandt (2020) showed that PLB tubules intersect to form tetrahedral units that are arranged in hexagons in their cryo-ET analysis of ruptured etioplasts, as expected from a zinc blende structure (*i.e*., diamond cubic lattice). We did not find any evidence for wurtzite-type lattices in our 3D models; this crystal type was detected in runner bean etioplasts in ET analysis by Kowalewska et al. (2016). The zinc blende lattice is a center-closest packed crystal system with a repeating unit of three layers, whereas the wurtzite lattice is a hexagonal closest packed system with a repeating unit of two layers (Cotton et al., 1995). We prepared approximately 300-nm thick sections to enclose more than four layers within PLBs. Due to the dark staining of stroma in cryofixed samples, tomograms of such thick sections captured in the brightfield TEM mode suffered poor signal-to-noise levels and they were not suitable for automatic segmentation. It was crucial to employ STET to enhance the membrane contrast of PLBs to determine their crystalline structure. We cannot rule out the existence of a wurtzite lattice as it was seen at the boundary between zinc blende crystal domains in squash etioplasts before (Murakami et al., 1985).

The LPOR–Pchlide–NADPH ternary complex binds to the lipid bilayer to produce membrane tubules *in vitro*, and the complex breaks apart upon illumination, mobilizing components required for constructing the photosynthetic membrane (Nguyen et al., 2021). We observed collapse of the crystalline order from the PLB periphery at 1 HAL, indicating that photo-activation of LPOR and subsequent Pchilde reduction begins in the PLB exterior. The loss of crystalline architecture at 1 HAL was characterized by randomized internodal distances, reduced branching per node, and thickening of tubules. The tetrahedral branching points were dislocated in the intermediate zone in 1 HAL PLBs. Lying between the inner crystalline and outer irregular regions, the intermediate zone is likely the sites in the PLB lattice in which LPOR–Pchlide–NADPH ternary complexes have disassembled immediately after light exposure.

One of the first events in the conversion of proplastids into chloroplasts is the formation of pre-granal thylakoids from tubule-vesicular thylakoid membranes (Liang et al., 2018). Our data indicate that pre-granal thylakoids develop from PLB tubules at the PLB-prothylakoid interface and CURT1A is involved in the transformation (Fig. 7A). Bloated thylakoids arose from *curt1a* PLBs and they failed to form grana stacks (Fig. 7B). CURT1A-GFP that rescued the *curt1a* phenotype concentrated to patches surrounding a PLB at 0 and 2 HAL. We speculate that these GFP-enriched sites are where pre-granal thylakoids and grana stacks are assembled (Fig. 7C). Decay of the crystalline PLB was slower in *curt1a*, probably due to the block in the transformation of PLB tubules into pre-granal thylakoids. All three *CURT1* isotypes, *1A*, *1B*, and *1C*, were transcriptionally active, and their gene products were detected in de-etiolating cotyledon specimens. However, *curt1b* and *curt1c* mutant lines did not exhibit defects in the PLB-to-pre-granal thylakoid transition.

**Figure 7.**
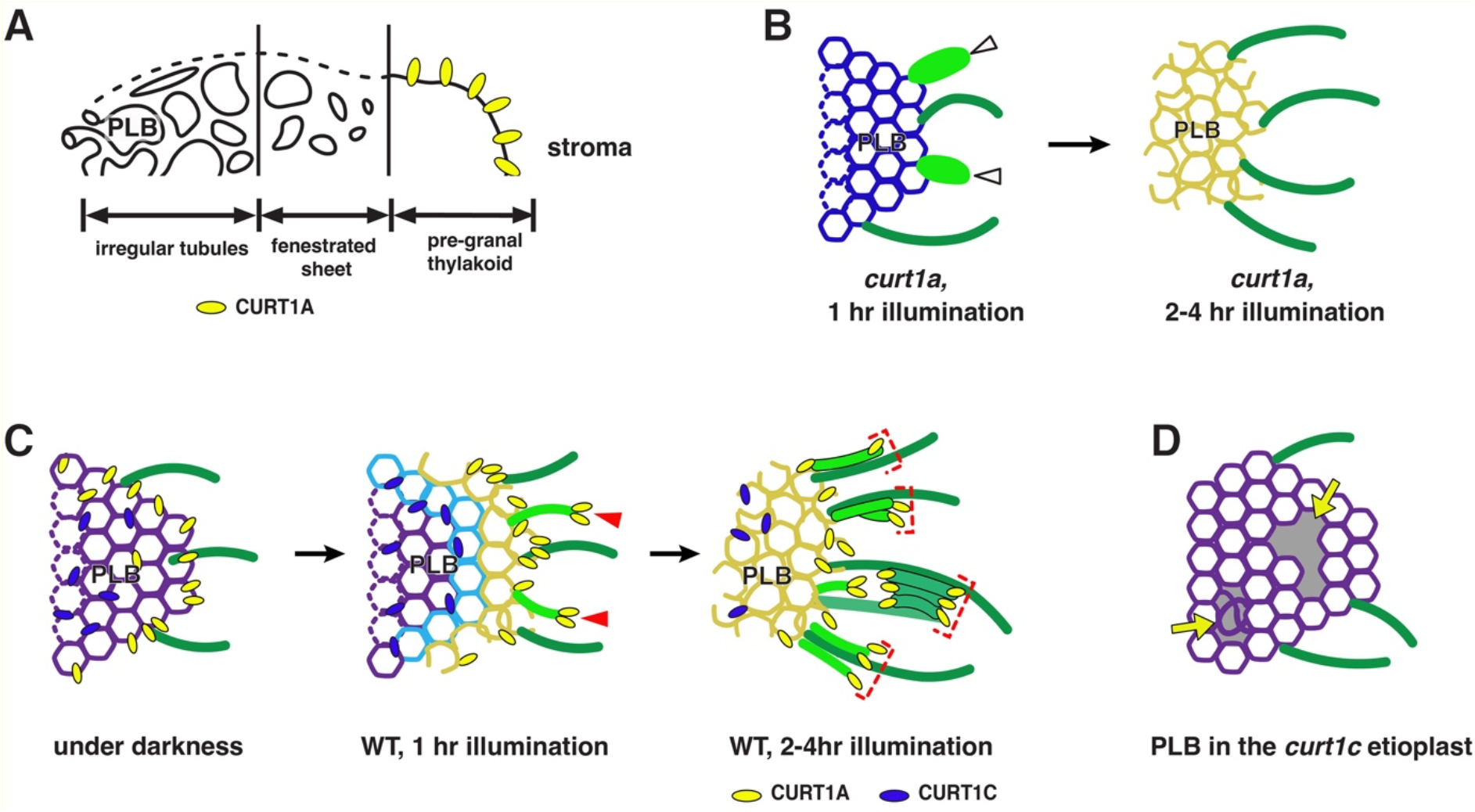
Schematic diagrams illustrating functions of CURT1A and CURT1C during deetiolation. (A) Model of pre-granal thylakoid development from PLB tubules. The irregular region of degrading PLBs gradually coalesced into a fenestrated sheet that matured into pregranal thylakoids when the fenestrae shrank to disappear. CURT1A stabilizes the membrane curvature at the tip of pre-granal thylakoids outgrowing from a PLB. (B) Pre-granal thylakoids growing out from PLBs are swollen (arrowheads) and grana stacks do not form in *curt1a*. (C) PLBs have CURT1A and CURT1C before light exposure. Upon illumination, PLB decay occurs from the margin, and CURT1A concentrates to the sites where new pre-granal thylakoids (arrowheads) and grana stacks (brackets) are assembled. CURT1C does not exhibit such relocation. (D) PLBs in *curt1c* mutant etioplasts have pores and disorganized tubules (arrows).

PLBs in *curt1c* mutant cotyledons had large holes or disarrayed tubules, indicating that CURT1C is required for PLB assembly in darkness(Fig. 7D). When we expressed CURT1C-GFP from the native *curt1c* promoter, the defects were rescued, validating a skotomorphogenic function of CURT1C. During de-etiolation, CURT1C-GFP spread uniformly over PLBs, and the PLB-associated fluorescence faded together with PLB degradation. The distinct mutant phenotypes and localization of GFP fusion proteins suggested that CURT1A interacts with the machinery for the production of pre-granal thylakoids in the PLB periphery. It will require 3D electron microscopic analyses of etiolating plastids at multiple time points after seedling germination under darkness to characterize the functions of CURT1C in PLB biogenesis.

In a recent publication, it was reported that the PLB structure and chloroplast biogenesis from etioplasts are affected in the *curt1abcd* quadruple mutant seedlings and that overexpression of *CURT1A* altered the PLB morphology (Sandoval-Ibanez et al., 2021). Our study is distinct from their research in that 1) our research is focused on the membrane dynamics involved in the conversion of PLB tubules into grana stacks occurring within 4 HAL, and 2) specified distinct functions of CURT1A and CURT1C (Fig. 7). Furthermore, we adopted high-pressure freezing for accurate 3D structural analysis of PLBs and membrane intermediates involved in the etioplast-to-chloroplast transformation. It is a consensus that organelle membranes, including the thylakoid, are preserved closer to their native states by high-pressure freezing than by conventional chemical fixation (Kiss et al., 1990; Kang, 2010; Nicolas et al., 2017; Otegui, 2021). Sandoval-Ibanez et al. (2021) preserved their EM samples by chemical fixation and 3 HAL was the first time point after 0 HAL when they examined PLBs and thylakoids using TEM/ET. Instead, they compared the photosynthetic capacity of *curt1abcd* and *CURT1A* overexpressor lines in more detail. Secondly, we characterized the functions of individual *CURT1* isotypes by examining their T-DNA mutant phenotypes and transgenic lines expressing GFP fusion proteins. It was impossible to uncover the roles of CURT1A and CURT1C separately in Sandoval-Ibanez et al. (2021) because their analyses were dependent on mutant lines lacking all CURT1 family proteins or overexpressing CURT1A. *curt1abcd* exhibited a delay in the onset of photosynthesis and changes in the PLB structure. These phenotypes of *curt1abcd* could be expected from defects in *curt1a* and *curt1c* single mutant lines (*i.e*., slower PLB-to-grana stack transition in *curt1a* and irregular PLB lattices in *curt1c*). However, the altered PLB structure in *curt1a* reported by Sandoval-Ibanez et al. (2021) does not match our result. PLBs in *curt1c*, not *curt1a*, exhibited abnormal crystal lattices. Furthermore, we did not see accelerated PLB degradation in *curt1a* etioplasts, unlike the quadruple mutant etioplasts. We could not interpret phenotypes of the CURT1A overexpressor line in the paper because the transgenic line was generated with Ler-0, of which PLB structure appears to be different from Col-0, and our study does not involve over-expressor lines.

## Methods

### Plant Materials and Growth Conditions

*Arabidopsis* Columbia (Col-0) and *curt1* seeds (NASC, http://arabidopsis.info/) were surface-sterilized and incubated in 4°C overnight. The seeds then were placed on 0.75% phytoagar Petri dishes supplemented with half-strength (0.5 g/L, pH 5.8) Murashige-Skoog salt (Sigma-Aldrich, USA; Cat. No. M5524). The dishes were placed in a growth chamber (Panasonic, Japan; Cat No. MLR-352H-PB) at 22°C and were left to germinate and grow for 1 week under darkness. Samples were harvested after illumination with white fluorescent light at a photon flux intensity of 120 μmol m^-2^ s^-^ before dissection.

### Generation of CURT1A-GFP, CURT1C-GFP lines in their respective mutant backgrounds

The genomic fragment of *CURT1A* (AT4G01150) and *CURT1C* (AT1G52220) including ~2 kb promoter region was amplified and inserted into a binary vector pBI121. The last exons of the genes were translationally fused with the GFP in the vector. *curt1a-1* and *curt1c-1* plants were transformed with the CURT1A-GFP and CURT1C-GFP constructs, respectively by floral dip method with the *Agrobacterium* tumefaciens strain GV3101(Zhang et al., 2006). Transgenic seedlings (T1) were selected by kanamycin containing 1/2MS + 0.8% phytoagar (w/v). Seedlings (T2 generation) were tested for GFP expression with immunoblot analysis (anti-GFP antibody, 1:2500 dilution, Abcam, USA; Cat. No. ab290) and observed under Leica TCS SP8 Confocal Microscope System (Leica Microsystems, Austria). All the primers were from Integrated DNA Technologies, and the genomic fragments were amplified with iProof high-fidelity DNA polymerase (Bio-Rad, USA; Cat. No. #1725301). Primer sequences for the GFP cloning are in Supplemental Table 2.

### High-pressure freezing, sample processing, and transmission electron microscopy

High-pressure freezing, freeze substitution, resin embedding, and ultramicrotomy were performed as described in Kang (Kang, 2010). Seedlings were examined with a Canon EOS M50 Digital Camera equipped with fluorescence illumination to remove abnormal cotyledons before freezing. Frozen samples were freeze-substituted in anhydrous acetone with 1% OsO4 at −80°C for 24 hr. Excess OsO4 was removed at −80°C by rinsing with precooled acetone. After being slowly warmed up to room temperature over 60 h, samples were separated from planchettes and embedded in Embed-812 resin (Electron Microscopy Sciences, USA; Cat. No. 14120). 80 nm thick sections of each time point were prepared with ultramicrotomy and then were examined with a Hitachi H-7650 TEM (Hitachi-High Technologies, Japan) operated at 80 kV.

### Dual-axis scanning transmission electron tomography, tomogram reconstruction, modeling, and measuring morphometric parameters

300 nm thick sections were collected on formvar-coated copper slot grids (Electron Microscopy Sciences, USA; Cat. No. GS2010-Cu) and stained with 2% uranyl acetate in 70% methanol followed by Reynold’s lead citrate (Mai et al., 2019). Tilt series from ±57° at 1.5° intervals in the scanning transmission electron microscopy (STEM) mode were collected with a 200-kV Tecnai F20 intermediate voltage electron microscope (Thermo-Fischer, USA). The FEI Tomography software (STEM mode) installed in the microscope was used to collect two tilt series around two orthogonal axes as described in Kang (Kang, 2016). Membrane surface models were generated according to the semi-automatic segmentation procedure in Mai and Kang (Mai and Kang, 2017).

### Immunoblot analysis and immunogold labeling

Protein samples were extracted from seedlings at 0 HAL, 1 HAL, 2 HAL, 4 HAL, 8 HAL and 12 HAL after being pulverized in liquid nitrogen. SDS-PAGE and immunoblot were performed as described by Liang et al. (2018) and Lee et al. (Lee et al., 2013). The experiment was repeated for three times with total protein extracts from three independent sets of cotyledon samples. For immunogold labeling, thin sections (80 nm thick) of HM20 embedded samples at each time point were prepared by ultramicrotomy, the following immunodetection of gold particles were performed according to the protocol explained in Wang et al. (Wang et al., 2017). Antibodies for PsaD (AS09 461), Lhca (AS01 005), AtpB (AS05 085), AtpC (AS08 312), PetC (AS08 330), PORA (AS05 067), CURT1A (AS08 316), CURT1B (AS19 4289), and CURT1C (AS19 4287) were purchased from Agrisera (Agrisera, Sweden). Anti-PBA1 antibody (ab98861) and anti-GFP antibody (ab6556) were purchased from Abcam (Abcam, USA). Antibodies against PsbO was provided by Michael Seibert (National Renewable Energy Laboratory). Antibodies for PsbP (Henry et al., 1997) and Lhcb (Payan and Cline, 1991) were donated by Kenneth Cline (University of Florida). For Fig S3C, multiple comparisons between data representing polypeptide readout of each time point and the readout of the 0 HAL using one-way ANOVA with Fisher’s LSD test. *p < 0.05, **p < 0.01, ***p < 0.001, ****p < 0.0001.

### Transcriptomic analyses

RNA samples were isolated from seedlings at each time point with 3 biological replicates using Qiagen Plant RNA extraction kit (Qiagen, Germany; Cat. No. 74904). A total of 18 cDNA libraries were prepared following the standard BGISEQ-500 RNA sample preparation protocol and sequenced by the DNBseq platform (BGI, China). Raw reads were filtered by SOAPnuke software, about 23.23 m clean reads for each sample were obtained in FASTQ format. The transcript expression level was then calculated and normalized to FPKM using RSEM software. The heat maps and the line charts were generated with R Studio (version 1.1.383) as described previously (Liang et al., 2018). FPKM values for CURT1 family genes were calculated to evaluate their expression level.

### Generation of skeleton models from PLB tubules

PLB membranes were first segmented using the 3D Orientation Field Transform tool (https://arxiv.org/abs/2010.01453). Skeletons were generated from the segmented membrane tubules by performing a medial axis transform (also known as ‘skeletonisation’) with an in-built MATLAB algorithm. Each skeleton element was converted into an undirected adjacency matrix carrying node coordinates using the Skel2Graph3D algorithm developed by (Kollmannsberger et al., 2017). The Bresenham’s line algorithm was used to connect node pairs with a straight line (https://arxiv.org/abs/2010.01453). The MATLAB adaptation of the Bresenham’s line algorithm iptui.intline() was modified for this purpose.

### Analysis of computer generated PLB skeleton models

The radial distribution function was computed by first plotting a histogram of the distances *r* between all the nodes in the skeleton, then the binned number was divided by *4πr^2^*. A curve approximating the histogram was used to generate probability plots against radial distances. Numbers of branches were counted from skeleton models at each time points. A distance transform with an in-built MATLAB algorithm was used on the binary segmented PLB tomograms to estimate PLB tubule thicknesses. The skeleton of the original segmentation was then introduced as a mask to select voxels around the central axes of PLB tubules. Approximate radii of PLB tubules were calculated from sizes of the voxels. The radii values were doubled to acquire diameters that correspond to tubular thicknesses. As we have calculated diameters from numerous voxels along auto-segmented PLB tubules, we were able to acquire low p-values (high degrees of confidence) in the pairwise comparisons.

## Accession Numbers

The RNA-seq data have been deposited in NCBI Sequence Read Archive under accession number GSE189497.

## Supplemental Data

The following material is available in the online version of this article.

Supplemental Figure 1. Etioplast-chloroplast transition in de-etiolating *Arabidopsis* cotyledons

Supplemental Figure 2. Generating skeleton models from PLB tubules in tomograms.

Supplemental Figure 3. Transcriptomic analysis and immunoblot analysis of proteins associated with the thylakoid membrane, CURT1 family proteins, and PORA.

Supplemental Figure 4. Characterization of *curt1* T-DNA inserted mutant lines.

Supplemental Figure 5. The abnormal thylakoid assembly phenotype reproduced in the *curt1a-2* (GK-805B04) allele and rescue of *curt1a* defects by expression of CURT1A-GFP

Supplemental Figure 6. Etioplast-to-chloroplast differentiation in *curt1b-1* cotyledons.

Supplemental Table 1. Statistics of the three rounds of RNA-seq experiments at the six time points.

Supplemental Table 2. Primer sequences for genotyping or molecular cloning

Supplemental dataset 1. The skeletal model file of a PLB shown in Figure 3.

## Acknowledgements

This work was supported by the Hong Kong Research Grant Council (GRF14121019, 14113921, AoE/M-05/12, C4002-17G) and Chinese University of Hong Kong (Direct Grants).

## Author Contributions

B-H. K. and Z.L. designed the research. Z.L., W-T.Y., J.M., K.K.M., Z.Y.L., Y-L. F.C., and X.C. performed the experiments. All authors analyzed the data. B-H. K. and Z.L. wrote the article.

## Supplementary Figures

**Supplemental Figure 1.**
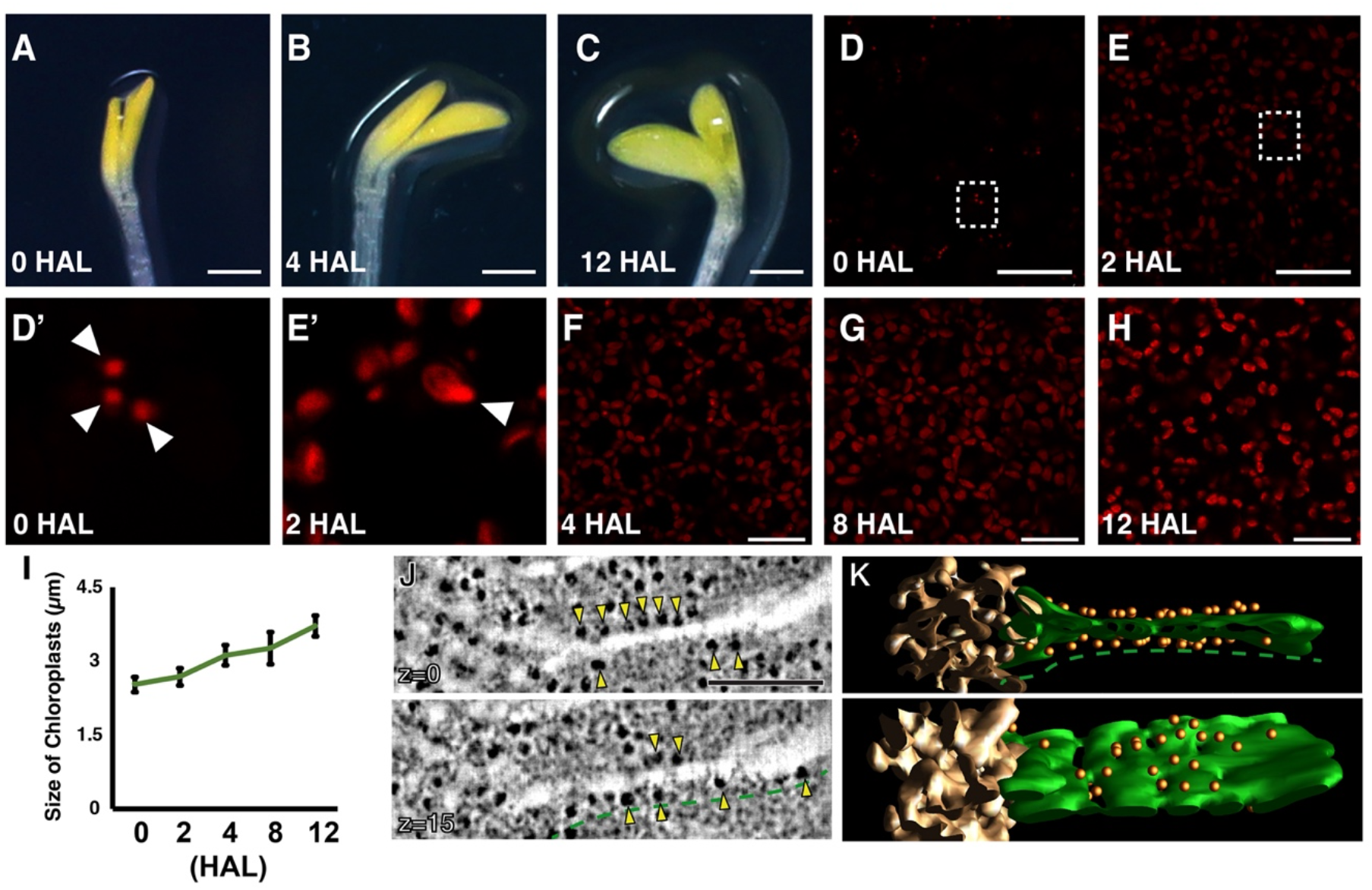
Etioplast-chloroplast transition in de-etiolating *Arabidopsis* cotyledons. (A-C) 7-day-old etiolated Arabidopsis cotyledons at A) 0 HAL, B) 4 HAL, and C) 12 HAL. Scale bars: 0.4 mm. (D-H) Confocal laser scanning micrographs showing Pchlide/chlorophyll autofluorescence at D) 0 HAL and E) 2 HAL., F) 4 HAL, G) 8 HAL, and H) 12 HAL. The excitation wavelength was 638 nm. Emission was detected in the wavelength range 651-715 nm. D’ and E’ show the areas marked by dashed squares in D and E, respectively. Arrowheads indicate PLBs in D’ and E’. Scale bars = 20 μm. (I) Increase in the plastid size during de-etiolation. Lengths of 30 plastids were measured in TEM images from three different cotyledon samples at each stage. Error bars indicate standard deviations. (J-K) STET slice images (J) and 3D models (K) of a prothylakoid (green) connected to a PLB (gold) at 0 HAL. The prothylakoid (green dash lines) has ribosomes (yellow arrowheads). Scale bars in (J): 200 nm.

**Supplemental Figure 2.**
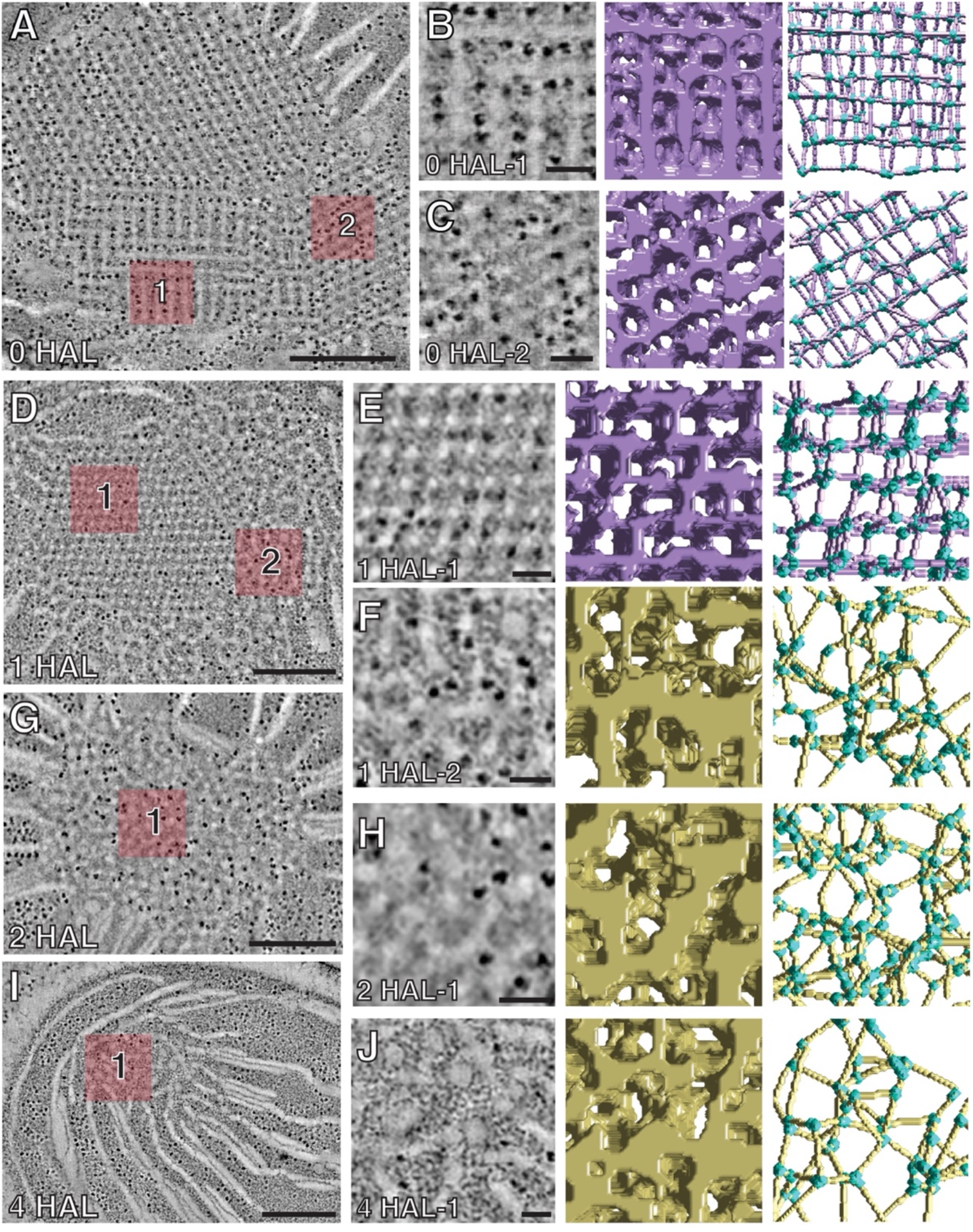
Generating skeleton models from PLB tubules in tomograms and prothylakoid membrane-bound ribosomes in the 0 HAL etioplast. (A-C) For a PLB at 0 HAL, panel A shows a STET slice image. In panels B and C, the numbered regions in panel A are magnified in the left panels, and automatic segmentation of PLB membranes and calculated skeleton models shown in middle and right panels. (D-F) For a PLB at 1 HAL, panel D shows a STET slice image. In panels E and F, the numbered regions in panel D are magnified in the left panels, and automatic segmentation of PLB membranes and calculated skeleton models are shown in middle and right panels. (G and H) For a PLB at 2 HAL, panel G shows a STET slice image. In panel H are shown a magnified view of the highlighted area in G (left), automatic segmentation of PLB membranes (middle), and calculated skeleton model (right). (I-J) For a PLB at 4 HAL, panel I shows a STET slice image. In panel J are shown a magnified view of the highlighted area in panel I (left), automatic segmentation of PLB membranes (middle), and calculated skeleton model (right). Nodes in the skeleton models are colored in blue. Scale bars in A, D, G, and I = 500 nm. Scale bars in B, C, E, F, H and J = 100 nm.

**Supplemental Figure 3.**
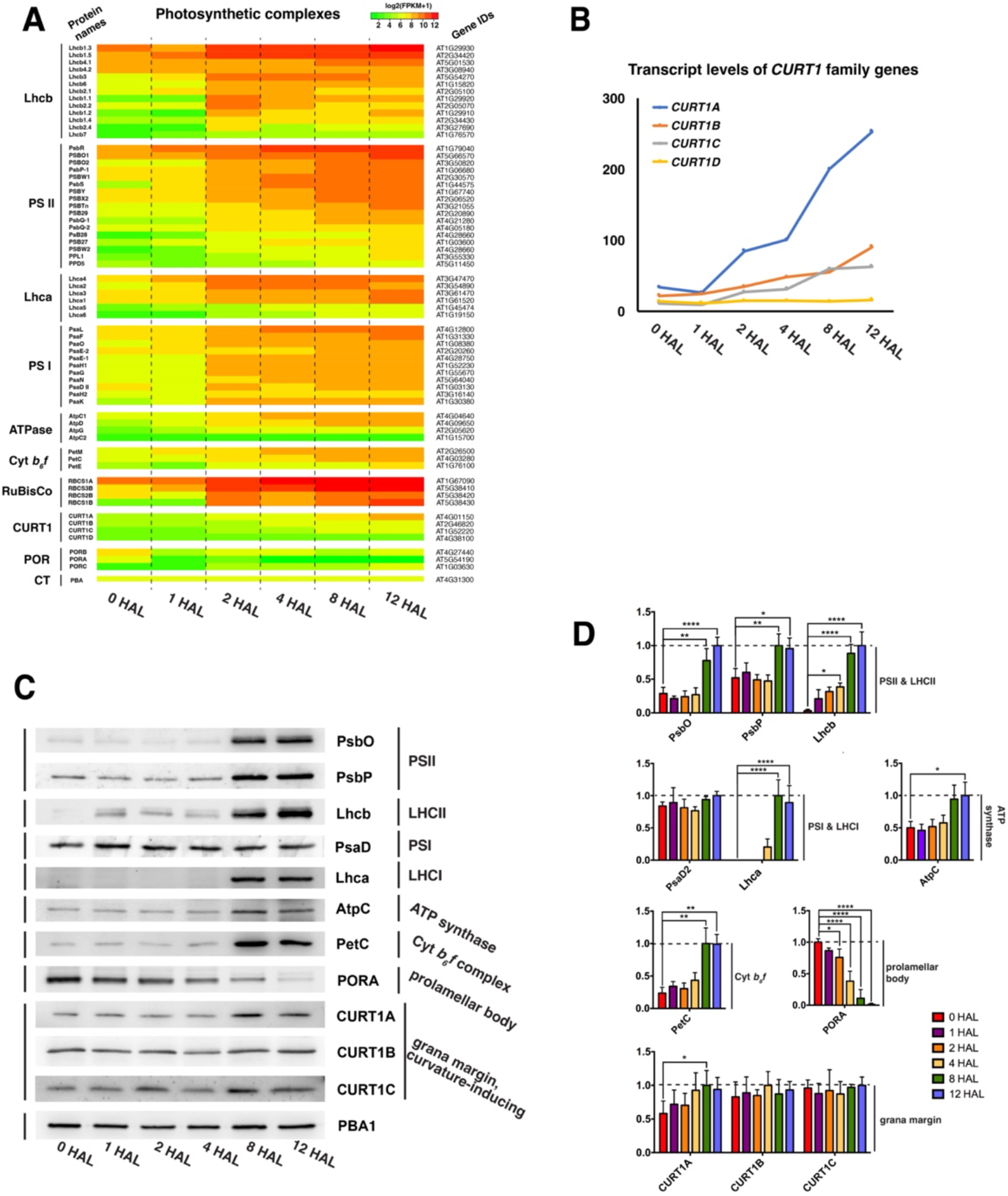
RNA-seq and immunoblot analyses of photosynthetic proteins, PORA, CURT1 proteins in de-etiolating *Arabidopsis* cotyledons. (A) Heat map illustrating levels of indicated transcripts in de-etiolating *Arabidopsis* cotyledons at six time points corresponding to those of microscopic analyses. Log2-fold changes are color-coded. Names of the subunits follow the nomenclature proposed by Race et al. (1999), and their gene IDs were obtained from www.arabidopsis.org. (B) Line charts showing levels of *CURT1* family transcripts. Note that transcriptional activity of *CURT1D* remained low in comparison to other *CURT1* genes. (C-D) Immunoblots of photosynthetic proteins, PORA, CURT1 proteins (C) and histograms showing their amounts quantified by chemiluminescence (D). The histograms were prepared with results from three repeats of immunoblot quantification. PBA1, a subunit of the Arabidopsis proteasome complex b, was used as the loading control. (±SD; one-way ANOVA; *p < 0.05, **p < 0.01, ***p < 0.001, ****p < 0.0001)

**Supplemental Figure 4.**
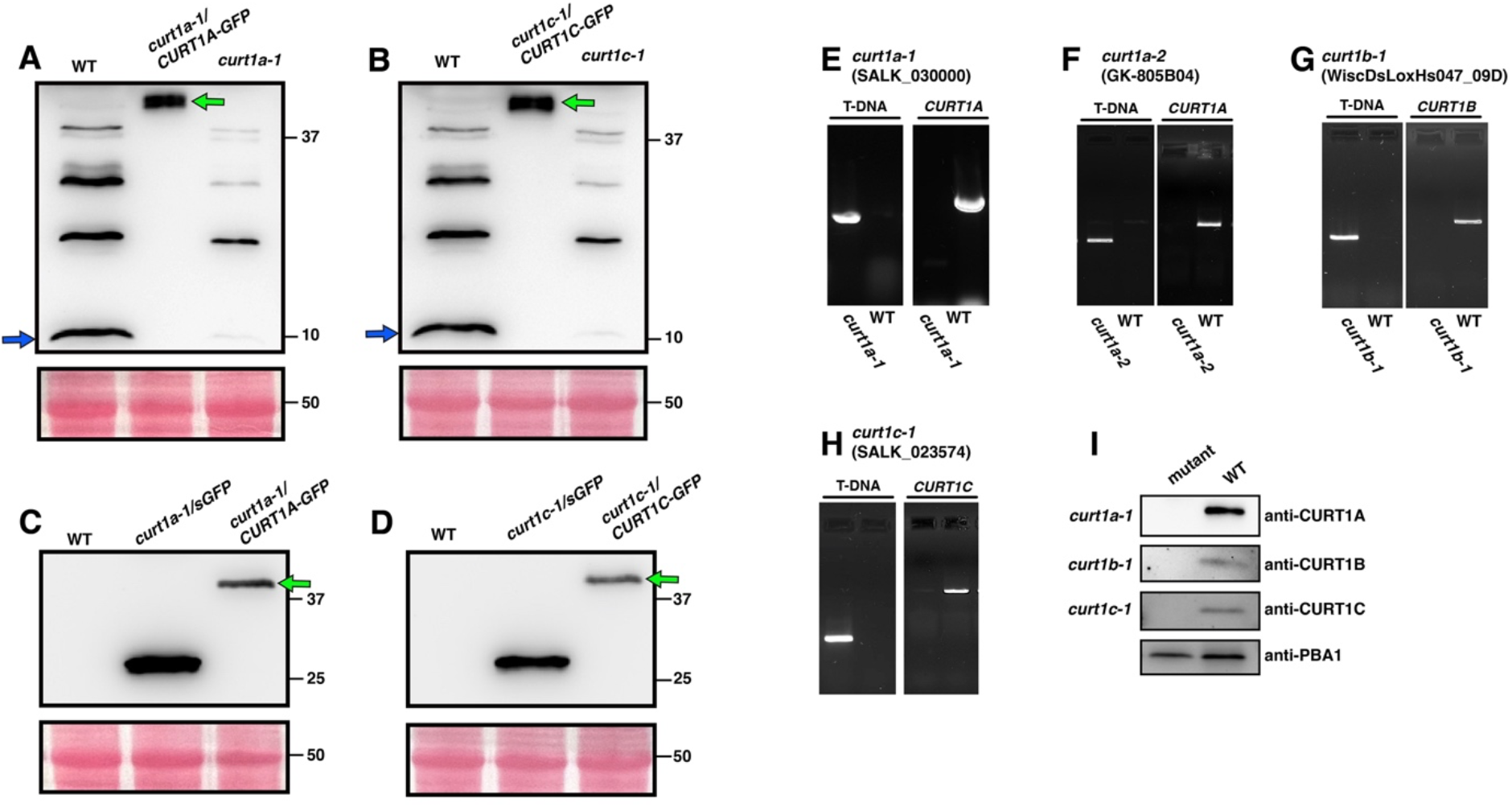
Characterization of *curt1* T-DNA inserted mutant lines. (A) Immunoblot analysis of wild type (WT), *curt1a-1* expressing GFP under its native promoters (*curt1a-1/CURT1A-GFP*) with an anti-CURT1A antibody. *(B)* Immunoblot analysis of WT, *curt1c-1* expressing GFP under its native promoter (*curt1c-1/CURT1C-GFP*) with an anti-CURT1C antibody. (C) Immunoblot analysis of WT, *curt1a-1/CURT1A-GFP*, *and curt1a-1* expressing a soluble GFP (*curt1a-1/sGFP) with a GFP antibody*. (D) Immunoblot analysis of WT, *curt1c-1/CURT1A-GFP*, *and curt1c-1/sGFP with a GFP antibody*. The mutant lines expressing sGFP were examined as positive control for GFP in (C-D). Ponceau S staining served as a loading control in (A-D). Blue arrows in (A) and (B) indicate CURT1A and CURT1C, respectively. Green arrows in (A-D) indicate CURT1-GFP fusion proteins. (E-H) PCR genotyping of intact and T-DNA-inserted *curt1a-1* (E), *curt1a-2* (F), *curt1b-1* (G), and *curt1c-1* (H) mutant lines. (I) Immunoblot analysis verifying the lack of CURT1A, 1B, and 1C proteins in their respective mutant lines. PBA 1 was detected as a loading control. Total protein extracts were prepared from 12 HAL WT and mutant cotyledon samples.

**Supplemental Figure 5.**
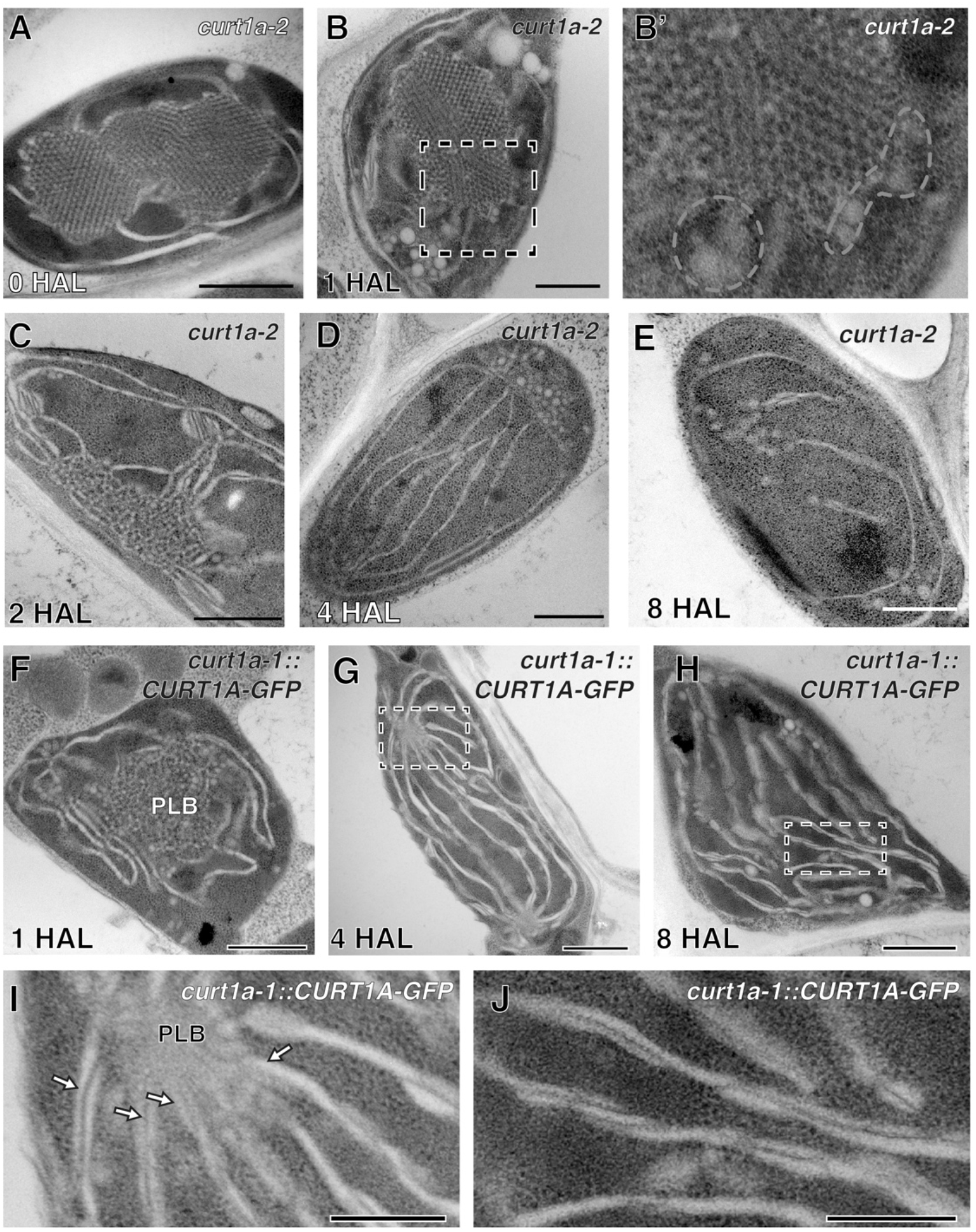
The abnormal thylakoid assembly phenotype reproduced in the *curt1a-2* (GK-805B04) allele and rescue of *curt1a* defects by expression of CURT1A-GFP. (A-E) TEM micrographs of the curt1a-2 at 0 HAL (A), 1 HAL (B), 2 HAL (C), 4 HAL (D), and 8 HAL (E). (B’) is the area enclosed in the dashed square in (B). Note the accumulation of swollen thylakoids around PLB in (B’). (F-H) TEM micrographs of etioplasts in *curt1a-1* expressing CURT1A-GFP at F) 1 HAL, G) 4 HAL, and H) 8 HAL. Panels I and J are higher magnification images of the area inside the dashed rectangles in G and H, respectively. Arrows indicate planar thylakoids emerging from PLBs in the GFP rescue lines. Scale bars in (A-H): 1.0 μm. Scale bars in (I) and (J): 200 nm

**Supplemental Figure 6.**
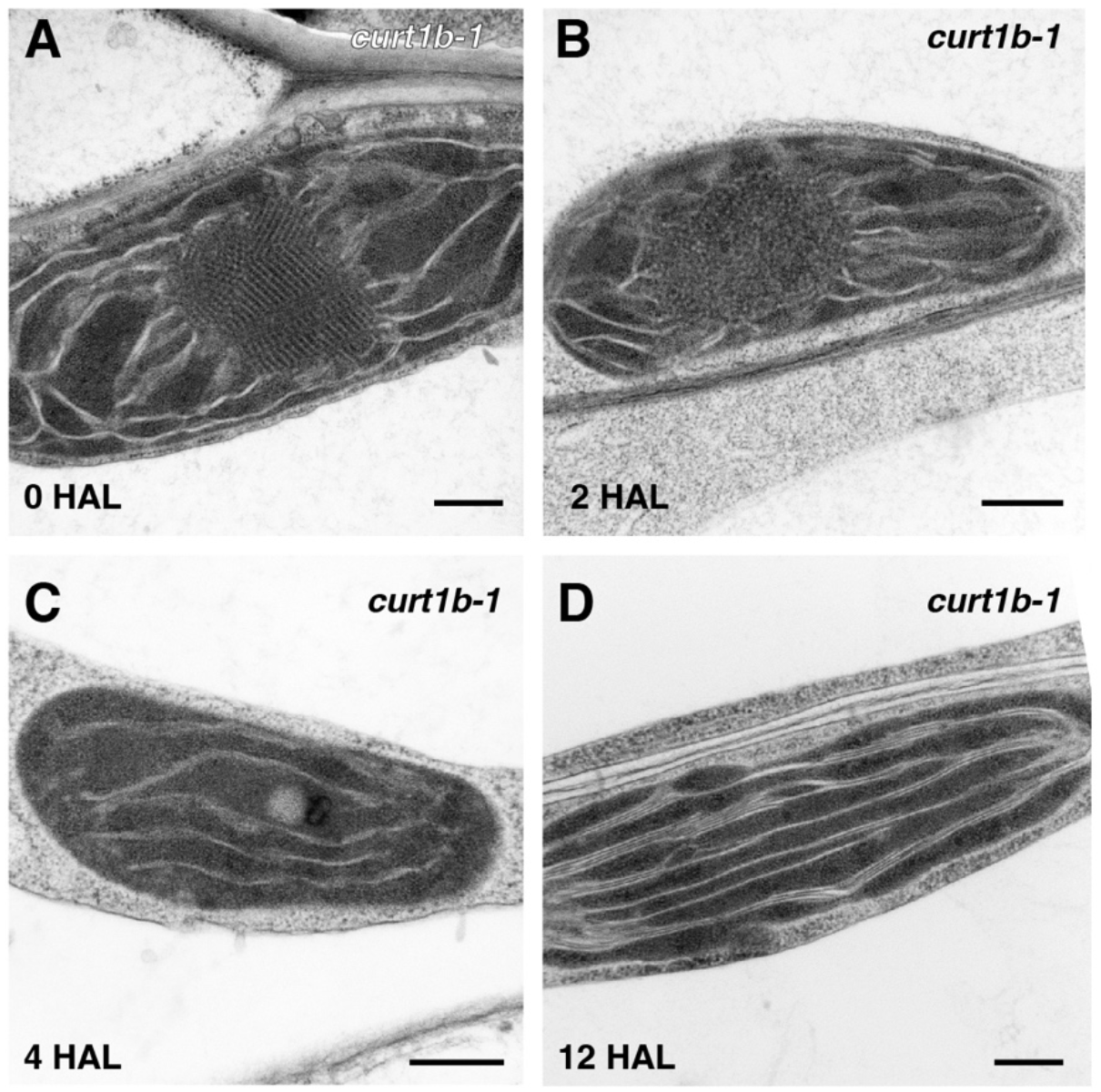
Etioplast-to-chloroplast differentiation in *curt1b-1* cotyledons. (A-D) TEM analysis of the *curt1b-1* (WiscDsLoxHs047_09D) plastids at 0 HAL(A), 2 HAL(B), 4 HAL(C), and 12 HAL(D). PLB degradation and grana assembly appeared normal in *curt1b-1* plastids (A-C). However, grana stacks in 12 HAL were extended and had less layers (D). Scale bars: 300 nm.

